# A Practical Refinement of Golgi Staining for Cortical Neuronal Morphology

**DOI:** 10.64898/2026.03.11.711075

**Authors:** Daniel Allen-Ross, Francesco Tamagnini, Maria Maiarú

## Abstract

Golgi-based staining remains a valuable method for visualising the full morphology of individual neurons, however conventional workflows can be slow, variable, and technically demanding. Here, we present a modified staining protocol designed to provide a practical workflow for Golgi-based labelling in mouse cortical tissue.

The revised workflow combines controlled fixation, dichromate impregnation, silver development, vibratome sectioning, and slide processing conditions that support visualisation of neuronal somata and dendritic architecture. In the tissue examined here, the protocol produced stained tissue with discernible neuronal morphology and was suitable for morphological assessment using standard light microscopy and image analysis software.

Overall, this method may be useful for laboratories seeking a straightforward and cost-conscious protocol for neuronal morphology.

## Introduction

Golgi-based staining has been a pivotal technique in neuroanatomy and neuroscience, uniquely enabling the visualisation of neuronal components, including the cell body (soma), basal and apical dendrites, and axons in their entirety. Discoveries made using Golgi staining techniques have fundamentally shaped our understanding of neuronal connectivity. Modern adaptations, including methodological refinements (Vints et al., 2019) and high pressure approaches (Tang et al., 2025) have positioned Golgi staining alongside advanced techniques such as genetic and fluorescent labelling within the neuroscientist’s toolkit, improving clarity, simplicity and cost-effectiveness (Rosoklija et al., 2014).

Historically, classic Golgi staining required prolonged incubation and impregnation, highly skilled operators, specialised equipment, and often produced inconsistent staining quality, including brittle tissue and high background artefacts. The development of modified protocols has aimed to reduce processing time, preserve morphology, improve accessibility, and minimise artefacts. These advances have sought to improve reproducibility, practicality, and processing efficiency. Importantly, such improvements enable high quality staining while reducing sample loss, which is especially important for studies of fine neuronal structures such as dendritic spines.

This workflow is intended for laboratories seeking clear visualisation of individual neurons for morphometrics analyses, spine quantification, soma size measurements, or neurocircuitry mapping across a range of tissue sizes, from rodent to non-human primate brain (Levine et al., 2013). It provides a practical alternative where advanced imaging or molecular approaches may be impractical or cost-prohibitive. By building on established histological principles, this protocol provides a practical resource for researchers aiming to visualise neuronal morphology in brain tissue.

The adapted Golgi-based staining protocol described here is intended to visualise neuronal cells, enabling detailed analysis of neuronal morphology, dendritic architecture, and connectivity in both research and diagnostic contexts (Wechsler et al., 1982). The aim is to provide a practical, accessible technique that addresses some practical limitations of classical Golgi methods in routine laboratory settings (Bayram-Weston et al., 2016; Du et al., 2023; Jiang et al., 2021; Pedrazzoli et al., 2021; Rosoklija et al., 2014; Zhang et al., 2020; Zhong et al., 2019). This methodology paper describes a methodological refinement and not a direct benchmark against competing protocols.

### Animal Ethics + Welfare

All animal procedures were conducted in accordance with the UK Animals (Scientific Procedures) Act 1986 and institutional ethical approval from the Home Office, under the appropriate project licence at the University of Reading. The study was carried out in compliance with relevant institutional and ARRIVE guidelines, and the 3R’s for the care and use of laboratory animals. Adult male mice, aged 20 weeks, were housed in groups of 5 per cage in a temperature-controlled facility maintained at 20 °C with a 12 h:12 h light/dark cycle. Animals had *ad libitum* access to food and water and were provided with nesting, millet, housing, tunnels and enrichment. Animals were acclimatised to the housing environment 7 days before experimentation and were monitored regularly throughout the study. All efforts were made to minimise animal suffering and to reduce the number of animals used, consistent with the principles of replacement, reduction, and refinement.

Only male animals were used because tissue was available from an ongoing study that also used male mice. This approach supports the 3R’s refinement principle by making use of tissue that was already available

### Methods (Protocol Section)

**Caution**: Potassium dichromate is toxic, carcinogenic, causes severe skin and eye burns, respiratory irritation, and may damage fertility and organs. Silver nitrate causes skin and eye irritation and lung irritation with repeated exposure. Both should be handled only in a properly functioning fume hood using protective gloves, goggles, and lab coats to avoid inhalation and contact hazards. Wash immediately and seek medical help if exposed

1. Preparation of Fresh Tissue

1.1 Deeply anaesthetise the mouse with 2.5% inhalation isoflurane in an induction box. And confirm death according to ASPA (1986) guidelines.

1.2 Excise the brain and immediately immerse it in 8ml of 10% formalin for 24 hours at 4 °C for a long term fixation process of tissue.

1.3 After 24 hours, discard the formalin and add 4 °C chilled 30% sucrose solution to act as a cryoprotectant and prevent ice crystal formation until the tissue is ready for slicing.

2. Golgi Staining

2.1 Discard the 30% sucrose solution and immerse the brain in 8ml of 3% potassium dichromate solution in a 15ml falcon tube. Protect the sample from light and incubate for 5 days at room temperature at 19°C, replacing the solution daily.

2.2 On Day 6, discard the potassium dichromate solution and add freshly prepared 2% silver nitrate solution kept at room temperature. Change the solution daily for another 5 days. **Note** that the solution will turn black due to a double displacement reaction forming insoluble silver dichromate, while potassium and nitrate ions remain in solution as spectator ions.

2.3 On Day 11, remove the stained brain tissue, blot gently, and place it into an appropriately sized well plate filled with 30% sucrose solution for vibratome sectioning.

3. Coronal Brain Slice Sectioning

3.1 Remove the brain from the sucrose solution, blot dry and orient it dorsally.

3.2 Slice off the cerebellum using a sharp blade, then cut along the sagittal plane to separate the hemispheres into left and right.

3.3 Apply cyanoacrylate adhesive to the vibratome stage and adhere the cerebellum side of the brain, ensuring that the olfactory bulb faces the blade.

3.4 Fill the vibratome stage with 30% sucrose solution and lock it into place.

3.4 Setup the vibratome and cut 60μm coronal slices at 0.12mm/s. Collect slices from the bath and transfer them to wells containing 30% sucrose solution.

4. Microscope Slide Application

4.1 Place left and right hemisphere sections onto microscope slide carefully to complete one coronal slice per animal. Ensure at least three technical replicates are adhered to the slide for each individual mouse.

4.2 Leave the slides for 3 hours to allow the brain slice sections to adhere before proceeding to the dehydration step.

5. Dehydration Step

5.1 Work inside a fume hood and align the adhered brain slices on the slides on absorbent paper.

5.2 Add one drop of 95% ethanol to each section and leave for 3 minutes. Invert the slide and allow excess solutions to drain onto the absorbent paper.

5.3 Add one drop of 100% ethanol to each section and leave for 3 minutes. Invert the slide and allow excess solutions to drain.

5.4 Add one drop of xylene to each section and leave for 2 minutes. Invert the slide and allow excess solutions to drain.

5.5 Apply one drop of mounting media to each section and carefully place a coverslip, ensuring no air bubbles are trapped around the brain slice.

6. Imaging

6.1 Examine Golgi-stained tissue sections under standard upright light microscopy (such as Zeiss Axioskope microscope fitted with GX Capture software). Capture images at 5x, 10x, 40x and 100x magnification.

6.2 Quantify cell body density within the region of interest (ROI).

### Reagents and Equipment

**Table 1.** List of materials needed through the golgi staining experiment stating material name, supplier and catalogue number.

| Reagent | Supplier | Catalog Number |
| --- | --- | --- |
| Formalin solution, neutral buffered, 10% | Sigma-Aldrich | HT501640-19L |
| MTC™ Bio Screw-cap Specimen Vial | Sigma-Aldrich | MTCV2250-700EA |
| Potassium Dichromate | Sigma-Aldrich | P5271-500G |
| Silver Nitrate | Sigma-Aldrich | 209139-25G |
| Sucrose | Sigma-Aldrich | S0389-1KG |
| Ethanol | Sigma-Aldrich | 51976-500ML-F |
| Xylene | Sigma-Aldrich | 247642-500ML |
| DPX Mountant for Histology | Sigma-Aldrich | 06522-100ML |
| Corning® microscope slides, frosted one side, one end | Sigma-Aldrich | CLS294875X25-72EA |

**Table 2.** List of equipment needed through the golgi staining experiment stating material name, supplier and catalogue number.

| Equipment | Model |
| --- | --- |
| Vibratome | Leica VT1200 |
| Upright Microscope | Zeiss Axioskope |
| GX-Capture | n/a |
| Fiji ImageJ | n/a |

### Representative Results

Application of the refined protocol produced stained 60 µm vibratome-cut brain slices suitable for morphological assessment and visualisation of neuronal morphology, including well-defined cell bodies and dendritic arbors suitable for morphometric analysis. In the initial study (see Fig 4), tissue from naïve mice (n=8) was processed using the refined protocol to assess somatosensory cortex tissue quality. Following successful staining, tissue sections were first imaged at 5x magnification, with higher magnifications subsequentially used to visualise individual neurons within the region of interest.

**Figure 1.**
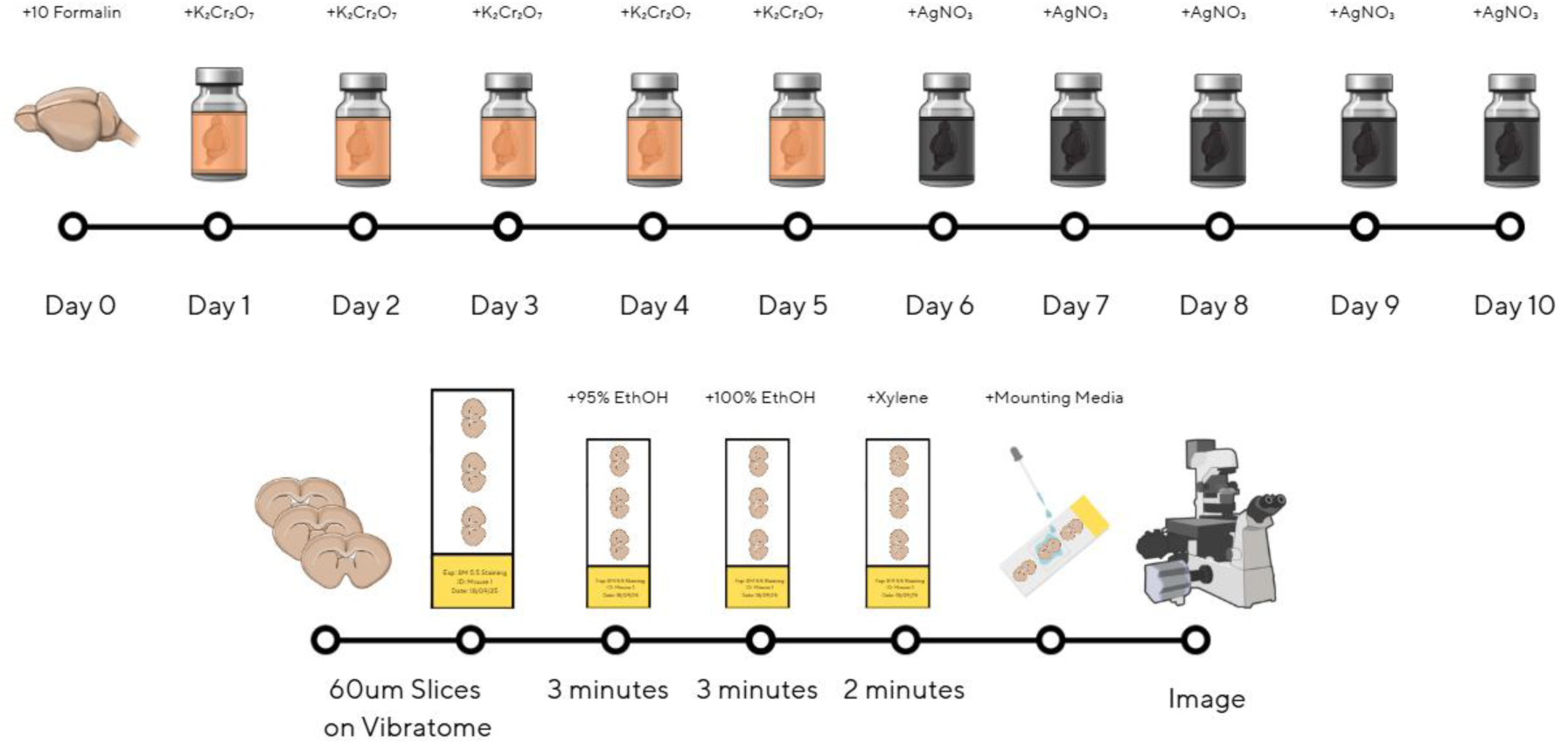
Schematic overview of the Golgi staining protocol for brain tissue slices. The upper panel illustrates the timeline and reagents used: brains are first fixed in 10% formalin for 24 hours, followed by sequential incubation in potassium dichromate (K₂Cr₂O₇), for five days and silver nitrate (AgNO₃) for an additional five days, with daily solution changes. The lower panel details the sectioning and processing steps: brains are sectioned into 60μm slices using a vibratome, then transferred to slides and processed sequentially in 95% ethanol (3 minutes), 100% ethanol (3 minutes), and xylene (2 minutes). Slices are then mounted with media before imaging for neuronal morphology quantification using an upright microscope and GX-Capture (Generated in Canva).

Figure 2 presents representative images derived from an experimental cohort of mice subjected to the SNI model and treated with psilocybin (4-phosphoryloxy-N,N-dimethyltryptamine) (Askey et al., 2025), a serotonergic psychedelic, compared with a saline control, using a standard Golgi staining technique. Power calculations and effect size were determined from previous experiments, indicating that n=4 was sufficient.

**Figure 2.**
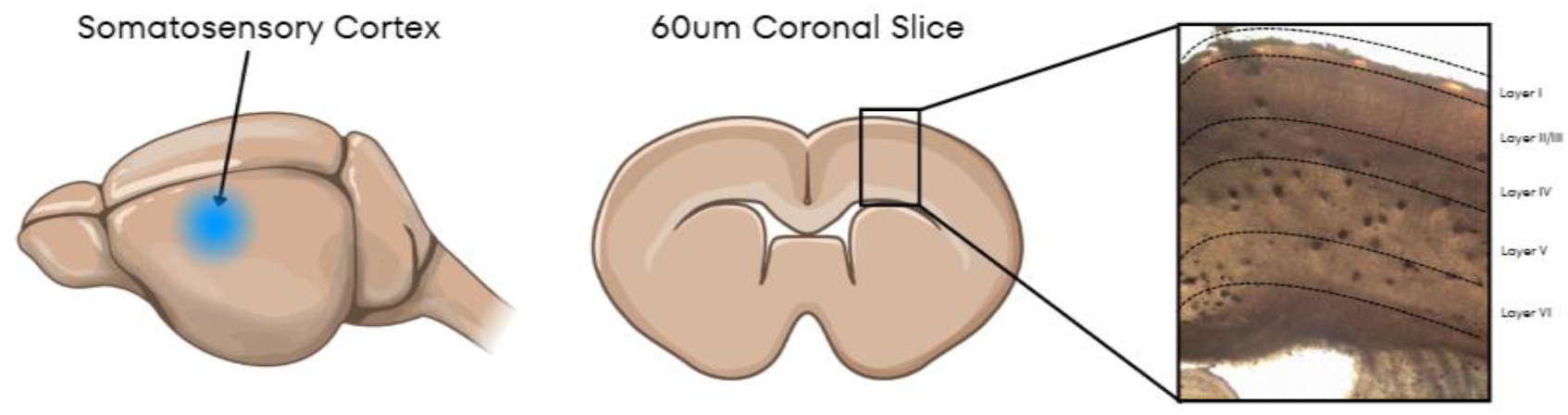
Schematic of the somatosensory cortex isolated in a single hemisphere where individual neuronal cell bodies through the brain layers have been golgi stained under a 5x Axioskope microscope lens, captured with GX-Capture at 60μm (Generated in Canva)

In a subsequent experimental study with further refinement (see Fig 3 and 5), tissue from SNI male mice (n=8) was processed to assess the integrity of somatosensory cortex structures. Following successful whole brain staining, three technical replicates per mouse were prepared to ensure consistency. Tissue sections were initially imaged at 5x magnification, followed by higher magnification imaging to examine individual neurons within the region of interest. Cell body density within the somatosensory cortex was then analysed and quantified to provide descriptive quantitative output.

**Figure 3.**
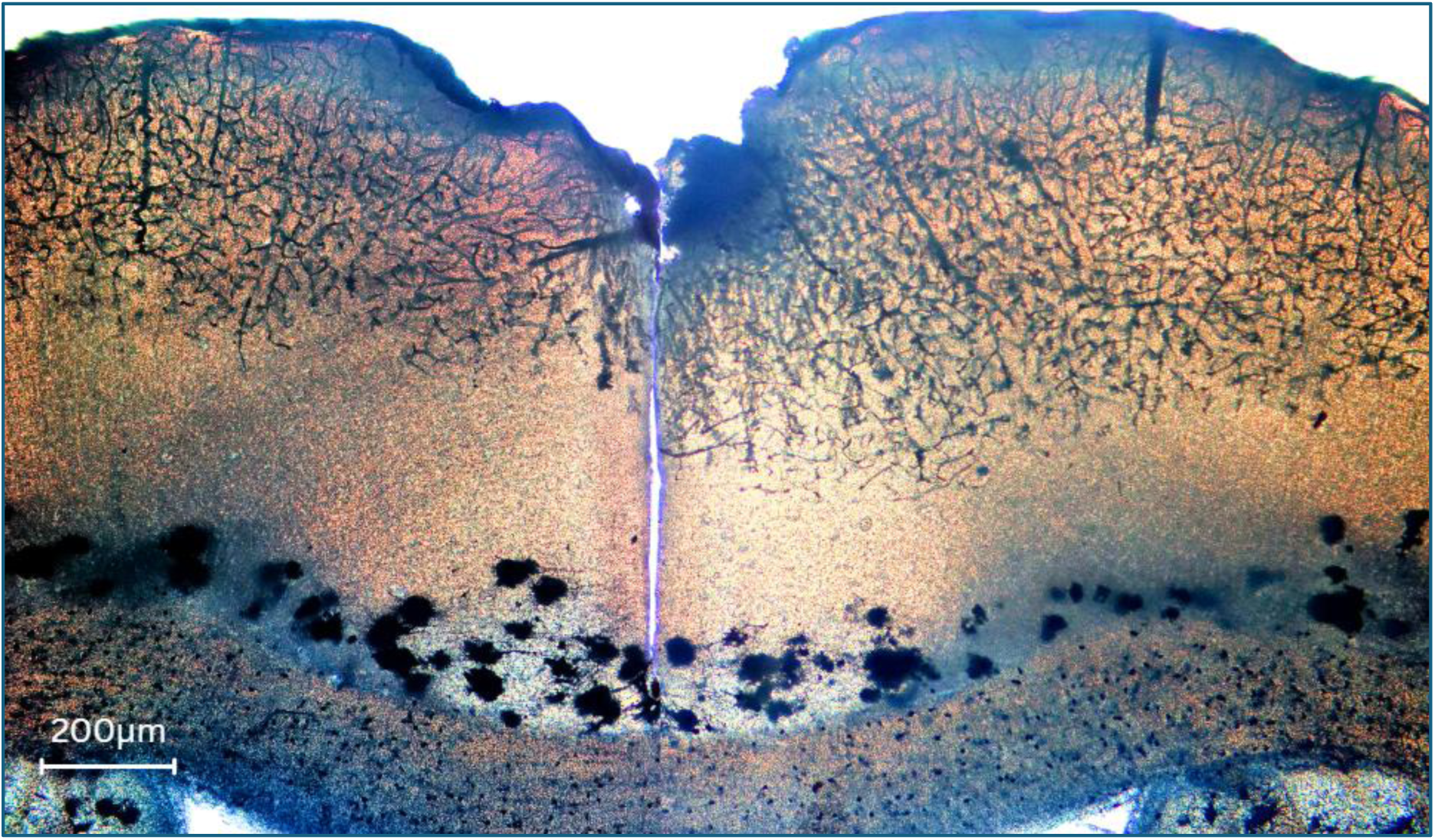
Representative image of a 60µm coronal mouse brain section stained using the refined Golgi-Cox method, visualised at 5x magnification. Neuronal cell bodies and dendritic structures are distinctly labelled throughout the cortex, showing discernible neuronal labelling suitable for morphological assessment. Scale bar, 200μm (Generated in Canva)

**Figure 4.**
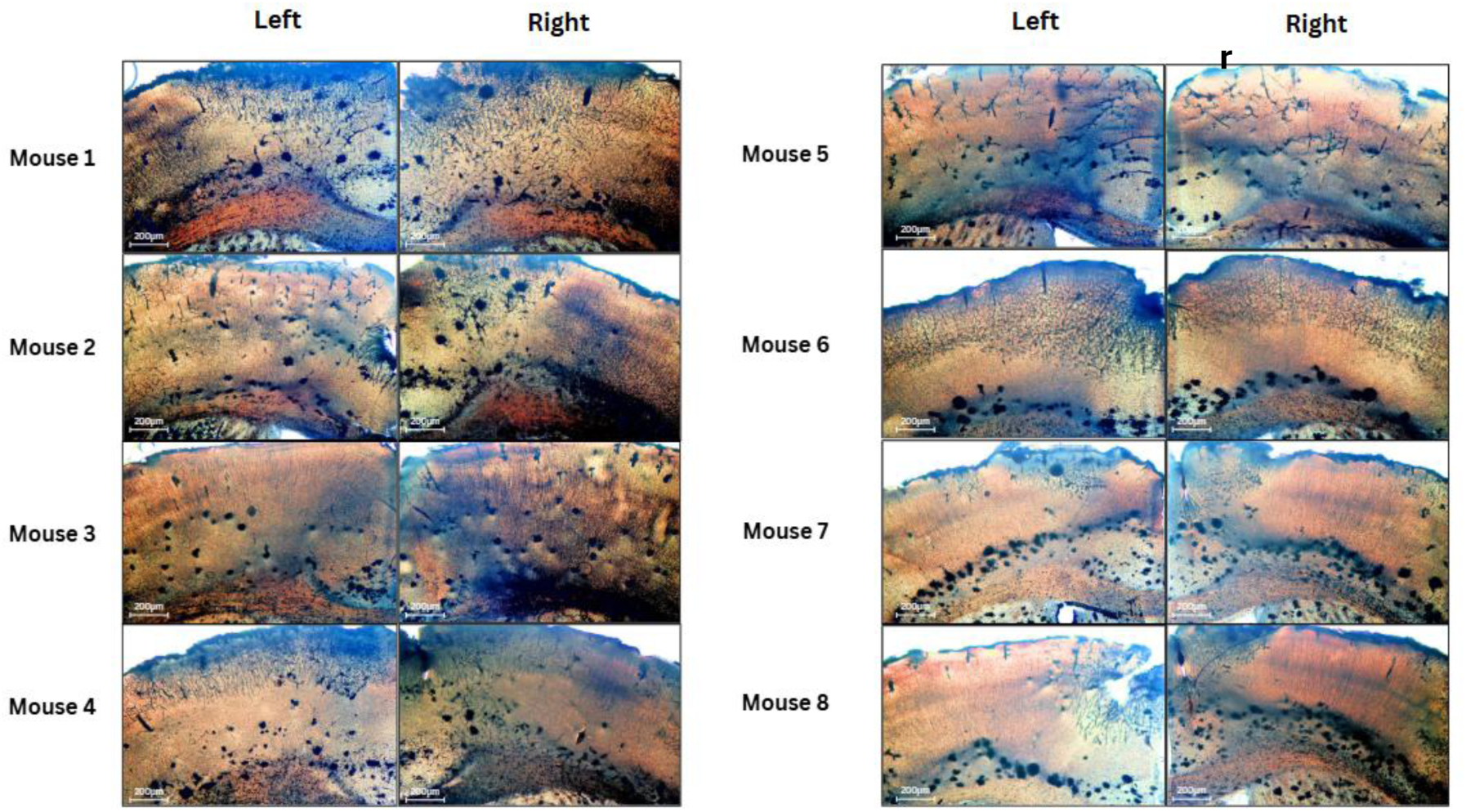
Representative 60µm coronal brain slices from 8 naïve mice processed with a refined Golgi-Cox staining protocol. For each mouse (Mouse 1 - Mouse 8), both left and right hemispheres are shown for comparative visualisation of staining consistency and neuronal morphology, labelling of cell bodies and dendritic structures is observed in multiple cortical layers and hippocampal regions, showing cellular detail that supports morphological visualisation. Scale bars, 200 µm (Generated in Canva).

**Figure 5.**
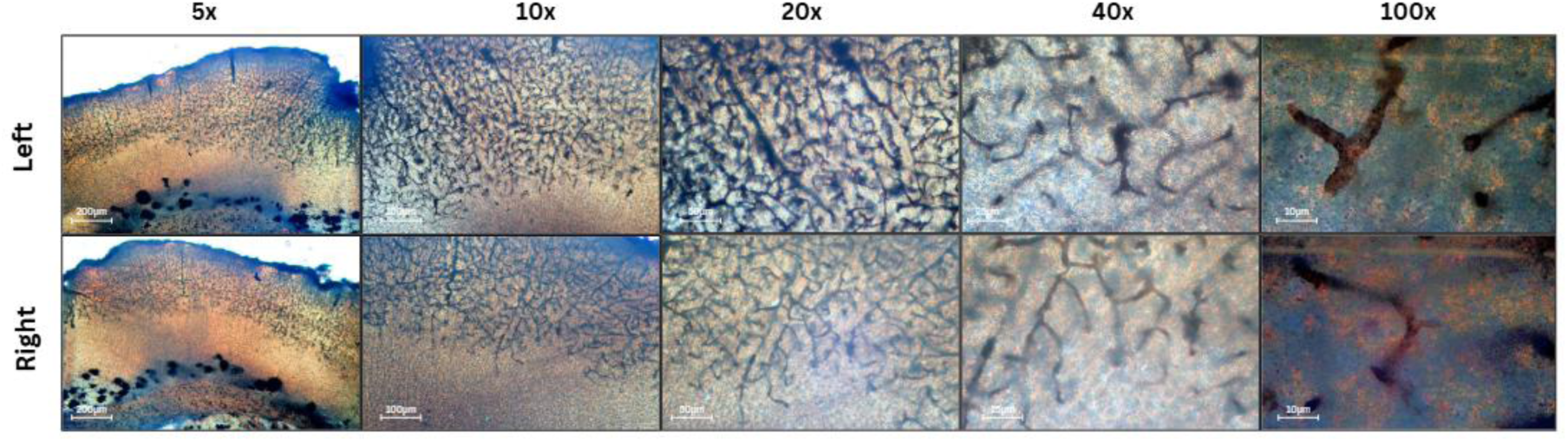
Representative images of mouse brain (Mouse 6) 60µm coronal sections stained with the Golgi method, visualised under increasing magnification. Slices from both the left and right hemispheres are shown in parallel rows for direct comparison. From left to right, columns display images captured at 5x, 10x, 20x, 40x, and 100x objectives. At lower magnification (5x, 10x), overall cortical architecture and distribution of Golgi-stained cells are visible. Progressively higher magnifications (20x, 40x, 100x) reveal increasing detail of neuronal dendritic arborizations and soma morphology, allowing visualisation of fine neural processes. Scale bars (200 µm for 5x and 10x, 50 µm for 20x, 20 µm for 40x and 100x) are included in each panel (Generated in Canva).

Collectively, these representative results indicate that the protocol can produce tissue suitable for neuronal morphology analysis under the conditions tested

### Statistical Validation

Cell body density was quantified in somatosensory cortex using a standardized region of interest (ROI) of 120,000 μm² in each hemisphere (left and right) per animal. For each mouse, three non-overlapping ROIs were sampled per hemisphere (3 technical replicates), and all clearly impregnated, non-overlapping somata fully contained within the ROI were counted. Raw counts were converted to areal density by dividing the number of cells by 120,000 μm² to obtain cells per μm² and then multiplying by 10⁶ to yield cell density in cells/mm² (cells per μm² × 1,000,000).

For statistical analysis, technical replicates were averaged within each hemisphere, and left/right hemisphere values were subsequently averaged to obtain a single cell density value per animal (n=8) and group data may be reported as mean ± SEM, where appropriate.

For dendrite/branch analysis in future studies, statistical approaches should be selected based on the specific measures (e.g. total branch length, branch order, Sholl intersections, branch points) and study design (n=4–5 animals/group, hemispheres, technical replicates), with animals treated as the primary unit of analysis to minimise pseudo-replication. For measures such as total dendrite length, branch points, or terminal endings, compute per-neuron means, average within hemispheres and animals, then apply one-way ANOVA (e.g, treatment: psilocybin vs saline) or two-sample t-test on group-level data.

For Sholl analysis (intersections vs radius): generate curves per neuron and perform a two-way repeated-measures ANOVA (radius as within-subjects, treatment as between-subjects) on animal-averaged data.

## Discussion

In this study, we describe a modified Golgi-based staining workflow for mouse cortical tissue that is intended to support practical usability and produces samples suitable for neuronal morphology assessment. The main contribution of the work is methodological: the protocol was refined to support visualisation of neuronal somata and dendritic processes under the conditions tested. The results therefore highlight the value of the method as a technical resource, rather than as evidence of a specific biological or mechanistic effect. Modifications such as fixation duration and impregnation temperature were adjusted to address common issues, including incomplete staining and background precipitate, thereby supporting clearer visualisation (Zaqout & Kaindl, 2016).

A major source of variation in Golgi staining protocols is staining impregnation durations. This protocol originally employed an 8 day staining period, consisting of 5 days in potassium dichromate, followed by 3 days in silver nitrate. Under these conditions, neuronal somata were stained, whereas dendritic morphology was less well resolved

Existing literature suggests that extending silver nitrate incubation can improve morphological definition. For example, Vints et al. (2019) reported a 15-day silver nitrate incubation for 100µm sections to enhance visualisation of hippocampal spines Mitchell et al. (2025) reduced a previous 16-day protocol to 11-12 days to optimise staining efficiency. Similarly, Vints et al. (2019) further extended impregnation to 22–36 days for thicker sections (150–500 µm), improving dendritic visualisation while minimising tissue loss. The present protocol, which visualises both somata and dendritic morphology, provides a foundation for further refinement within the conditions tested.

Conventional Golgi staining often fails at the impregnation step, producing few labelled neurons, pale staining, and high background noise, which can obscure fine structures such as dendritic spines. Louth et al. (2017) demonstrated that optimisation of impregnation conditions can substantially improve staining consistency, indicating that such artefacts are methodological rather than unavoidable. Common sources of failure include insufficient or excessive impregnation time, degraded or poorly prepared staining solutions, and suboptimal tissue handling. Practical consideration for optimising staining include:

- **Weak staining**: extend impregnation duration, refresh and filter staining solutions, and review reagent concentrations.
- **Patchy staining**: reduce tissue thickness, ensure adequate solution volume, and maintain stable, dark storage for the tissue.
- **High background noise**: slightly shorten impregnation duration and refine dehydration and drying step.

Despite these improvement, limitations remain inherent to the Golgi method. There is still no clear mechanism of staining between the reagents and the tissue being inherently stained. The stochastic nature of neuronal impregnation means only a small fraction of neurons (approximately 5% or fewer) are labelled per tissue sample, which complicates comprehensive sampling and may introduce bias in quantification (Koyama, 2013). The method is labour intensive and slower compared to modern fluorescent genetic labelling, often requiring multiple days to weeks for optimal impregnation depending on tissue type and thickness (Mitchell et al., 2025; Zaqout & Kaindl, 2016). Additionally, Golgi staining lacks molecular specificity and cannot discriminate neuron subtypes without additional labelling, limiting its use when specific neuronal populations are the focus (Rosoklija et al., 2014) and a generic quantification is required.

Compared with alternative techniques such as fluorescent protein expression or viral labelling, the Golgi method uniquely provides comprehensive, high-resolution labelling of entire neuromorphology including cell body soma, dendritic spines and fine processes, without requiring transgenic models or specialised microscopy (Zhang et al., 2020). This makes it valuable for studying non-model species or human postmortem tissue. Furthermore, recent protocol iterations have improved staining speed and tissue preservation, enabling compatibility with immunostaining and transgenic mouse models, broadening its versatility and utility in contemporary neuroscience research (Zhang et al., 2020), (Hui et al., 2025)

Our protocol has many advantages over existing techniques currently conducted by researchers. This refined methodology includes several practical features relative to classical Golgi-based methods compared classical Golgi-based methods (Risher et al., 2014) or marketed rapid staining kits in delivering clear labelling of neuronal components, enabling morphological analysis. This modified protocol is designed to reduce sample brittleness, and to shorten staining time relative to some longer conventional workflows (Zhang et al., 2020). Staining times are shortened, enabling results to be obtained within a relatively short period, allowing for a faster turnout for experimental work. There is no need for external thermal sources (Hui et al., 2025) or external pressure sources (Tang et al., 2025). Artefacts and background staining were reduced under the conditions used, which can improve the interpretability and quantification of neuronal components in the region of interest. The extended, shortened staining duration (See Fig 3 and 5) enhances impregnation quality, supporting visualisation of dendritic structures, including spines of cortical morphology with the potential to use dendritic spine tracking software. Lastly, the workflow uses standard histological reagents and equipment that avoids reliance on expensive reagents, high specification equipment or a highly specialised skill set by an individual.

Ranjan and Mallick (2012) also show that Golgi staining can be substantially improved by modifying incubation conditions, including temperature and fixation exposure which can lead to faster and more reliable labelling in rat tissue. However, those protocols were designed around different tissue processing conditions, species and readouts than the present mouse cortical workflow so differences in staining intensity, background noise and morphology are expected. Accordingly, our study should be interpreted as a refined protocol of cortical staining rather than a direct comparison with other Golgi refinements (See Table 3).

A major strength of the revised workflow is its simplicity. By using standard histological reagents and a straightforward sectioning and mounting procedure, the method avoids reliance on specialised commercial kits or advanced imaging systems. This makes the protocol attractive for laboratories that need an accessible approach for morphology-based studies. The present data suggest that the protocol can yield tissue suitable for downstream quantification of neuronal structure, including soma density and dendritic morphology, provided that processing is carried out carefully and consistently.

At the same time, the manuscript should be interpreted within clear limits. This study does not establish superiority over all existing Golgi, Golgi-Cox, or commercial rapid-staining protocols, because direct side-by-side benchmarking was not performed. The conclusions are therefore restricted to the performance of the adapted workflow described here. Likewise, although the stained tissue is suitable for structural analysis, the present data do not support broader claims about plasticity, functional circuit remodelling, or disease-specific effects. Those topics require dedicated experimental designs and independent validation. The study is limited by sample size and by the use of tissue from male animals only. Finally, this workflow may be a useful practical resource for laboratories seeking a straightforward approach to cortical neuronal morphology.

**Table 3.** Comparison of representative Golgi-based staining approaches for neuronal morphology analysis for neuronal morphology analysis. This table summarises the general chemistry, strengths, limitations, and most common applications of the major Golgi family staining approaches discussed in the revised manuscript. The present study is positioned as an optimisation of the protocol described in this paper, rather than as a direct experimental comparison

| Protocol type | Core chemistry / kit | Typical strengths | Typical limitations | Best use case |
| --- | --- | --- | --- | --- |
| Classical Golgi | Potassium dichromate and silver nitrate impregnation of fixed tissue | Simple conceptually, historically foundational, can reveal full neuronal morphology including soma, dendrites, and spines. | Highly variable impregnation, technically sensitive, often inconsistent background and section quality. | General neuron morphology when a classic stain is acceptable and protocol optimisation is available. |
| Golgi-Cox | Chromate-based impregnation with mercuric chloride in standard Golgi-Cox workflows | Strong neuronal labelling, widely used for dendritic and spine morphology, often good structural clarity. | Longer processing, protocol sensitivity, toxicity concerns due to mercury-containing chemistry | Detailed dendritic and spine analysis, especially in established morphology workflows |
| Commercial rapid-staining kits | Proprietary kit-based Golgi-Cox variants such as FD Rapid GolgiStain | More standardized, user-friendly, often faster and easier to implement across | Costly, kit dependence, still may need optimisation for tissue type and section thickness | Labs prioritizing convenience, reproducibility, and reduced setup complexity |

**Table 4.** Comparison of the present refined Golgi workflow with conventional or prior Golgi-based approaches. The table highlights the principal procedural features, and the scope of the current study. The present work should be interpreted as a protocol refinement rather than a direct comparative benchmark.

| Feature | Conventional / prior Golgi workflow | Present refined workflow | Rationale |
| --- | --- | --- | --- |
| Tissue fixation | Fixation conditions vary across laboratories and are not always specified in detail. | Brain fixed in 10% formalin for 24 h at 4 °C before impregnation. | Standardised fixation step to improve procedural consistency. |
| Impregnation chemistry | Classical Golgi and Golgi-Cox methods often require prolonged impregnation and are sensitive to handling. | Sequential potassium dichromate (5 d) followed by silver nitrate (5 d), with daily solution changes. | Defined incubation schedule intended to support consistent impregnation. |
| Sectioning | Section thickness and orientation are often protocol-dependent. | Vibratome sectioning at 60 µm with explicit orientation steps. | Provides a repeatable sectioning workflow for cortical tissue. |
| Slide processing | Dehydration and mounting steps are sometimes reported briefly. | Explicit 95% ethanol, 100% ethanol, and xylene sequence with defined incubation times. | Improves transparency and reproducibility of slide processing. |
| Scope | Some workflows are tailored to specific tissue types or specialised equipment. | Designed as an accessible cortical workflow using standard histology reagents and equipment. | Intended to be practical for routine laboratory use. |
| Comparative evidence | Direct benchmarking is not always included. | No head-to-head comparison with alternative Golgi methods was performed in the present study. | Limits conclusions to the workflow described here. |

## Acknowledgments

We thank Prof Gary Stephens for their constructive review and insightful feedback on this manuscript.

## Author contributions

DAR and MM conceived and designed the experiments; DAR performed the experiments; DAL, FT and MM drafted the paper and DAR wrote the manuscript. All authors revised and edited the manuscript.

## Data and materials availability

All data associated with this study are available in the main text.

## Supplementary Information

**Supplementary Table S1. ARRIVE 2.0 compliance statement**

This study has been reported in accordance with the ARRIVE 2.0 guidelines. The manuscript addresses the ARRIVE Essential 10, including study design, sample size, inclusion and exclusion criteria, randomisation where applicable, blinding where applicable, outcome measures, statistical methods, experimental animals, experimental procedures, and results

### Raw Images

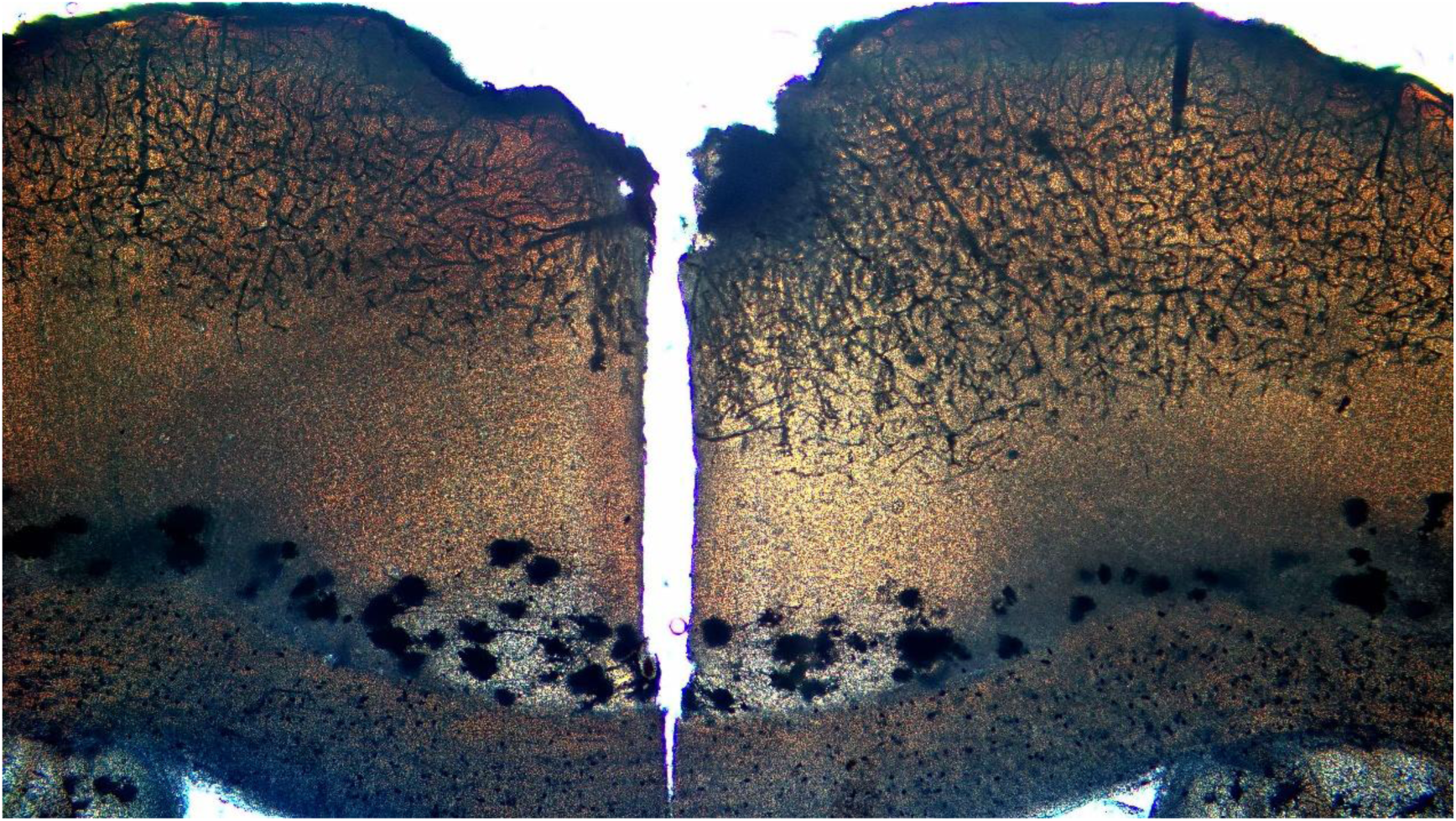

Raw image of a 60µm coronal mouse brain section stained using the refined Golgi-Cox method, visualised at 5x magnification. Neuronal cell bodies and dendritic structures are distinctly labelled throughout the cortex, demonstrating clear impregnation and high contrast suitable for quantitative morphology analysis. Scale bar, 200μm (Generated in Canva)

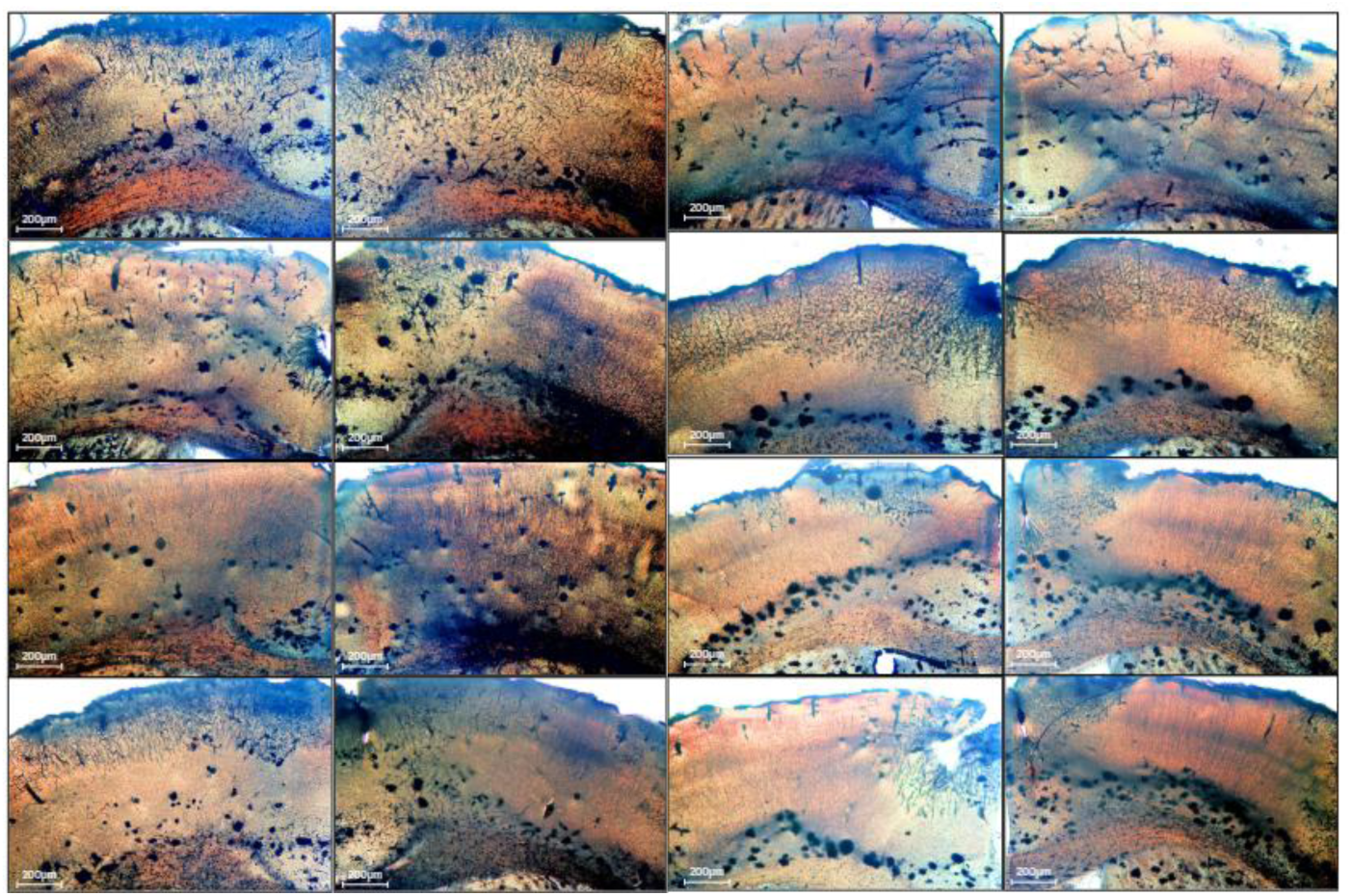

Raw images of 60µm coronal brain slices from 8 naïve mice processed with a refined Golgi-Cox staining protocol. For each mouse (Mouse 1 - Mouse 8), both left and right hemispheres are shown for comparative visualisation of staining consistency and neuronal morphology. Clear impregnation of cell bodies and dendritic structures is observed in multiple cortical layers and hippocampal regions, demonstrating clear cellular detail suitable for quantitative morphological analysis. Scale bars, 200 µm (Generated in Canva).

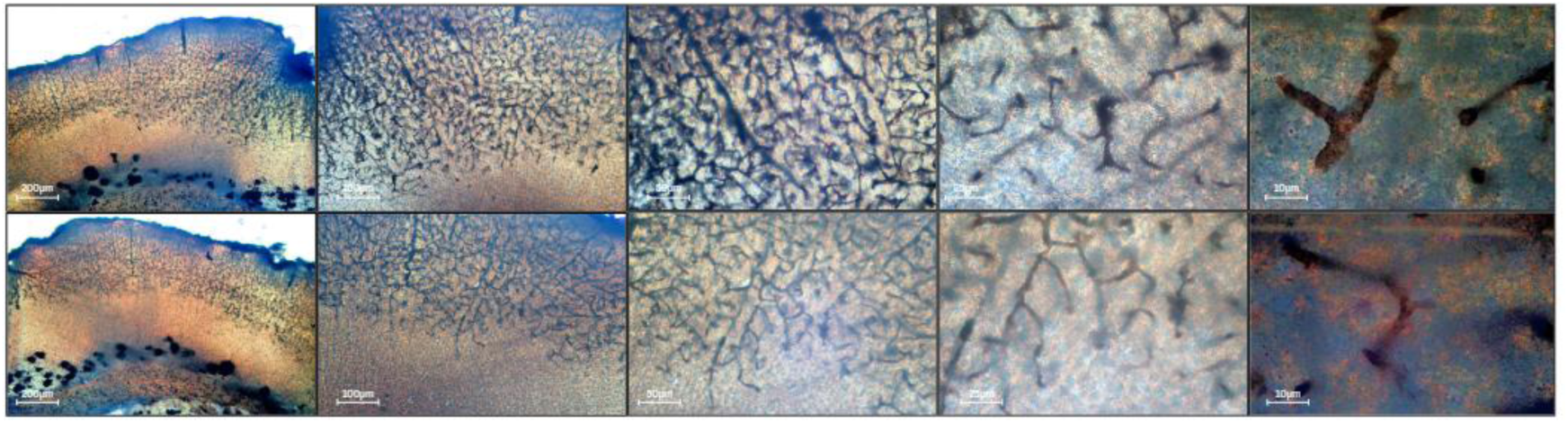

Raw images of mouse brain (Mouse 6) 60µm coronal sections stained with the Golgi method, visualised under increasing magnification. Slices from both the left and right hemispheres are shown in parallel rows for direct comparison. From left to right, columns display images captured at 5x, 10x, 20x, 40x, and 100x objectives. At lower magnification (5x, 10x), overall cortical architecture and distribution of Golgi-stained cells are visible. Progressively higher magnifications (20x, 40x, 100x) reveal increasing detail of neuronal dendritic arborisations and soma morphology, showing clear staining and enabling the visualisation of fine neural processes. Scale bars (200 µm for 5x and 10x, 50 µm for 20x, 20 µm for 40x and 100x) are included in each panel (Generated in Canva).

